# Therapeutic potential of methylxanthine drug on alpha synuclein protein in Parkinson’s disease

**DOI:** 10.1101/2021.12.20.473414

**Authors:** Nishant Kumar Rana, Neha Srivastava, Bhupendra Kumar, Abhishek Pathak, Vijay Nath Mishra

## Abstract

Parkinson’s disease (PD) is the second most common neurodegenerative disorder after Alzheimer. It exists in sporadic (90 to 95%) and familial (5 to 10%) form. Its pathogenesis is due to oxidative stress, glutamate excitotoxicity, protein aggregation, neuroinflammation and neurodegeneration. There is currently no cure for this disease. The protein-protein interaction and gene ontology/functional enrichment analysis have been performed to find out the prominent interactor protein and shared common biological pathways, especially PD pathway. Further *in silico* docking analysis was performed on target protein to investigate the prominent drug molecule for PD. Through computational molecular virtual screening of small molecules from selected twelve natural compounds, and among these compounds methylxanthine was shown to be prominent inhibitor to SNCA protein that ultimately prevent PD. The interaction of methylxanthine compound with the target protein SNCA suggested that, it interacted with prominent binding site with good docking score and might be involved in blocking the binding of neuroinducing substances like: 1-methyl-4-phenyl-1,2,3,6-tetrahydropyridine (MPTP) to SNCA protein. Thus methylxanthine compounds can be explored as promising drugs for the prevention of Parkinson’s disease.

## Introduction

Parkinson’s disease (PD) is a common neurodegenerative disorder with a prevalence of 160/100 000 in Western Europe and is rising to ∼4% of the population over 80 [1]. In most people, the symptoms appear at 60 years of age or beyond. A crude occurrence rate of 14.1 per 100,000 individuals was reported from rural Kashmir in the northern belt of India. The incidence rate of PD over the age of 60 years was 247 per 100,000 individuals [2] while a low incident rate was reported in south India i.e. 27 per 100,000 individuals in Bangalore [3]. In eastern part of India, 16.1 individuals per 100,000 populations were affected from PD [4] while maximum incidence ration (328.3 per 100,000 individuals) was observed in western India [5]. Early-onset Parkinson’s disease (EOPD) accounts for 3 to 5% of all PD cases while in other cases it is classified either into juvenile (below age 21) or ‘young-onset’ PD (age range of 21-40) [6]. The pathological hallmark of PD is cell loss within the substantia nigra particularly affecting the ventral component of the pars compacta. By the time of death, this region of the brain has lost 50–70% of its neurons compared with the same region in unaffected individuals [7]. Although PD is usually a sporadic disease, there are a growing number of single gene mutations which have been identified. Total 11 genes have been mapped by genetic linkage with six genes identified: α-synuclein (SNCA), ubiquitin C-terminal hydrolase like 1 (UCH-L1), parkin (PRKN), LRRK 2, PINK 1 and DJ-1 genes [8]. Parkin (PRKN) gene mutations are highest among all other genes reported in PD and vary from 1.96% to 39.1% among Indian PD patients [9]. DJ1, LRRK2 and PINK1 mutations are less frequent in PD while mutations in SNCA are absent [10]. Pathways concerned to PD pathogenesis include alpha-synuclein pathobiology, neuroinflammation, impaired protein degradation, synaptic and mitochondrial dysfunction [11]. There is a lack of uniform, large scale, nationwide epidemiological data on the incidence and prevalence of PD in India [12]. A survey done in Indian population shows the prevalence rate of PD to be 33-48 cases per 100,000 [13]. However, the Parsi community in Mumbai showed a prevalence rate of PD around 192 per 100,000, which was higher compared to rest of the Indian population [14]. Element imbalance in biofluids such as serum and CSF has been used for the development of diagnostic marker of PD [15]. Studies have reported several biomarkers in other diseases or in other types of biological fluids which are being pursued as blood-based biomarkers in PD [16].

Formerly coffee has been prohibited in many diseases, but now the negative views changed fundamentally; as the data shows that the Methylxanthine (MTX) exhibit health benefits in diseases related to nervous system involving cell death [17-18]. Caffeine, theophylline and theobromine are the mainly recognized members of MTX compounds; they are primarily present in tea, coffee, cacao, cola drinks and yerba mate. MTX is mainly absorbed in the gastrointestinal tract and go through into the central nervous system, where they exert significant psychostimulant actions [18]. Present study summarized the therapeutic potential of MTX in PD.

Designing of drugs that can target the protein involved in neurodegeneration may be an effective approach for the treatment of PD [19]. Identification of the neurodegeneration mechanisms from these natural agents has shed light on where they interact with the target protein [20]. Through computational molecular virtual screening of small molecules from natural compounds, have been confirmed to directly inhibit these important proteins to prevent the PD.

## Methods

### Protein-protein interaction (PPI) network analysis

The search tool for retrieval of interacting proteins (STRING) (https://string-db.org) database, which integrates both known and predicted PPIs, can be applied to predict functional interactions of proteins [21]. To analyze the interaction among selected proteins we have submitted the protein name on String database and retrieved the possible interaction among these proteins.

### Pathway analysis and network modeling

The molecular interaction network of selected proteins was created in Cytoscape v3.2.1 tool using proteins-biological pathway data set. Cytoscape network modeling tool determines the function of set of proteins or subgraph of a biological network. The main advantage of networking over other gene ontology (GO) tools is the fact that it can be used directly and interactively on molecular interaction graphs [22]. The interaction of selected proteins has been designed on the basis of shared common biological pathways. In order to find out the interaction among proteins, data were submitted in excel file format including the candidate proteins-pathway or biological process to identify their interaction and network. Further network has been generated on the basis of shared common biological pathways by using submitted data in excel sheet format as well as online search using several online databases.

### Retrieval of herbal compounds as a ligand

The 2D structure of herbal compound as a ligand were obtained from the PubChem Compound database (https://pubchem.ncbi.nlm.nih.gov/), followed by its conversion from the SDF to PDB format. Further the drug likeness property of herbal compound was checked by Lipinski filter [23].

### Molecular docking of SNCA protein with selected herbal compounds

Docking of ligands/drug molecules with various selected targets was carried out by PatchDock Server [24] (default parameter RMSD esteem 4.0 and complex type protein-small ligand) further visualization of the docked complexes through Discovery Studio 3.5. Docking investigation was based on geometric shape complementarily score (GSC score), which was determined in PatchDock Server, The findings of the results are based on the docking score and the interaction at the binding sites (pockets). The higher scores represents more binding affinity, and hence more steady is the complex. The docking assessment was done using Discovery Studio 3.0 to find out the contacting receptor residues involved in interaction with ligands. [25].

## Results

### Protein Protein interaction network analysis fot the selected proteins

Interaction among proteins which involved in neurodegeneration diseases, mainly PD has been performed using STRING Database. In this networking all proteins which involved in neurodegeneration, mainly PD such as: APP, SNCA, STUB1, LRRK2, PARK1, PARK2, PARK7, SNCAIP, FYN, HSPA4, UCHL1, and SLC6A3 were connected with each others. Among these interacted proteins the SNCA was the prominent interactor (Fig 1).

**Fig. 1.**
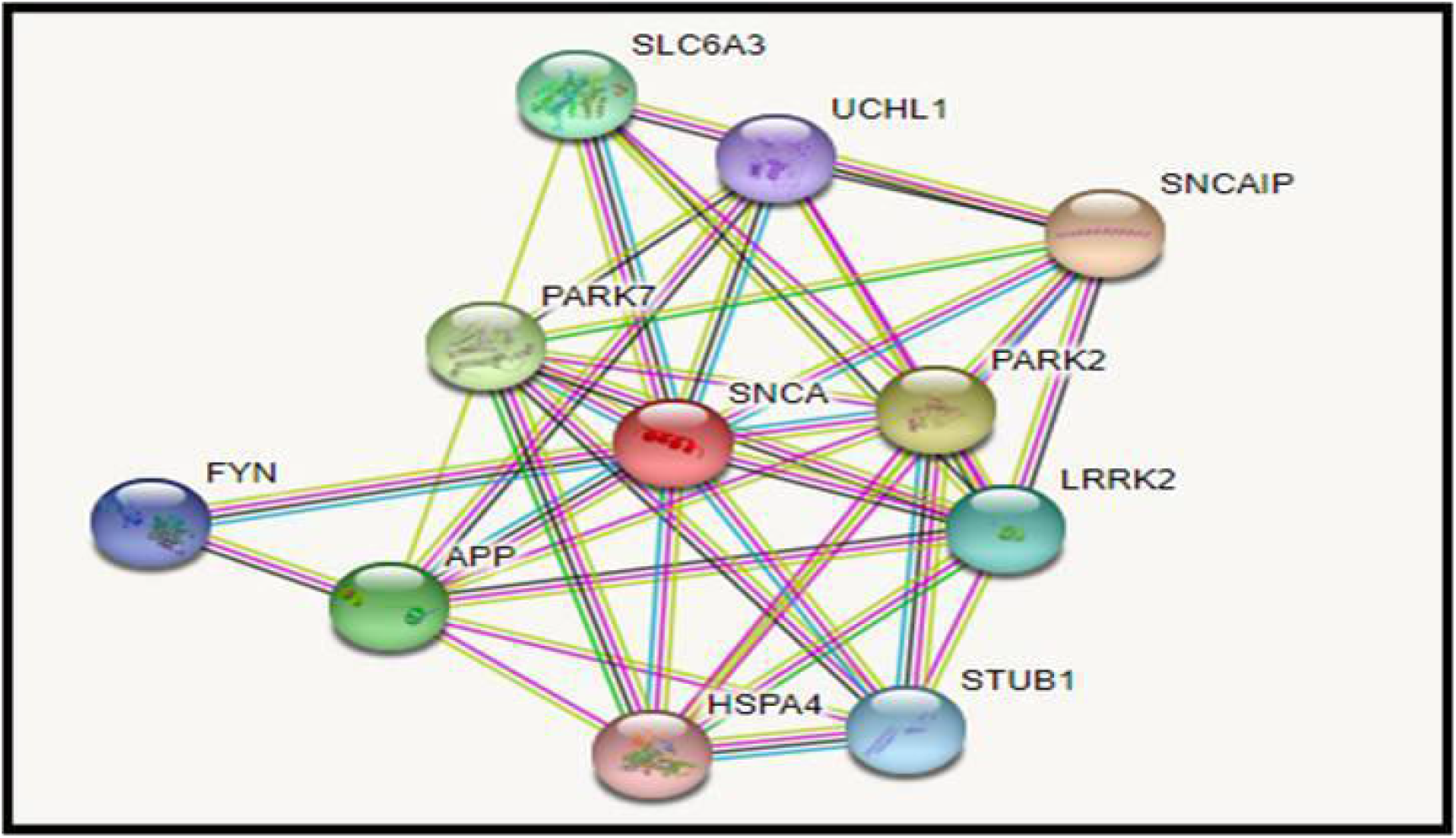
Network of protein-protein interaction involved in neurodegeneration

### Biological pathway analysis and network modeling

The network was generated in Cytoscape plug-in using targeted proteins association data set with their biological pathways. All experimentally validated target proteins or some other target proteins involved in Parkinson disease were retrieved from the experimental data set and create the network on the basis of shared common biological pathways. Networking of selected proteins such as: APP, SNCA, STUB1, LRRK2, PARK1, SNCAIP, FYN, HSPA4, UCHL1, and SLC6A3 were performed on the basis of their shared common biological pathways. From this network, it has been found that the SNCA protein was more prominent interactor and involved in cell neurodegeneration to cause PD (Table 1) (Fig 2).

**Table 1.**
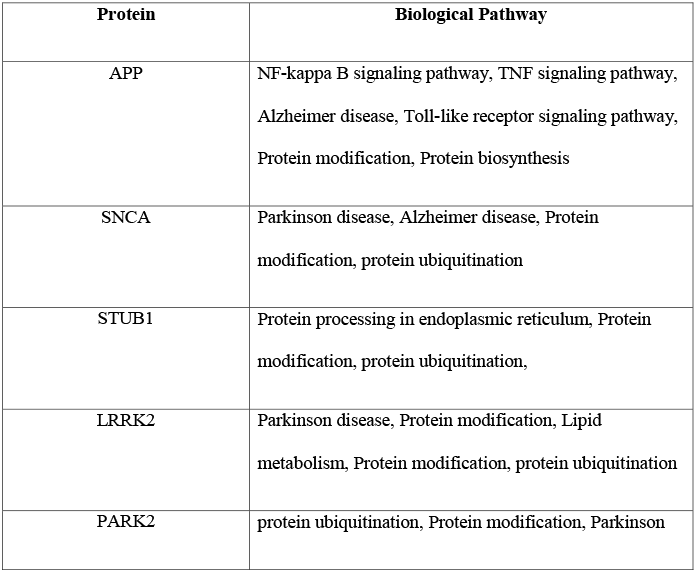
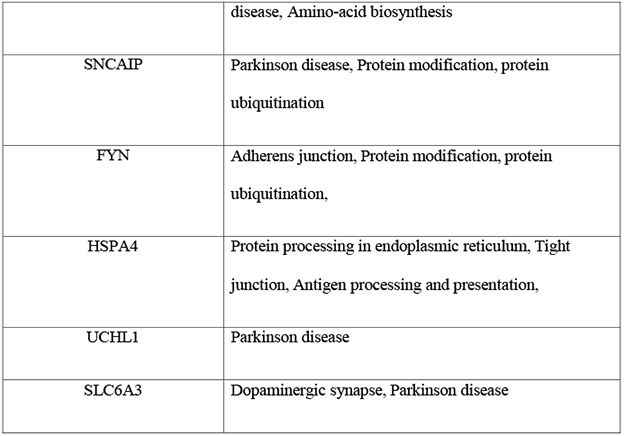
GO analysis of selected proteins involved in neurodegeneration

**Fig. 2.**
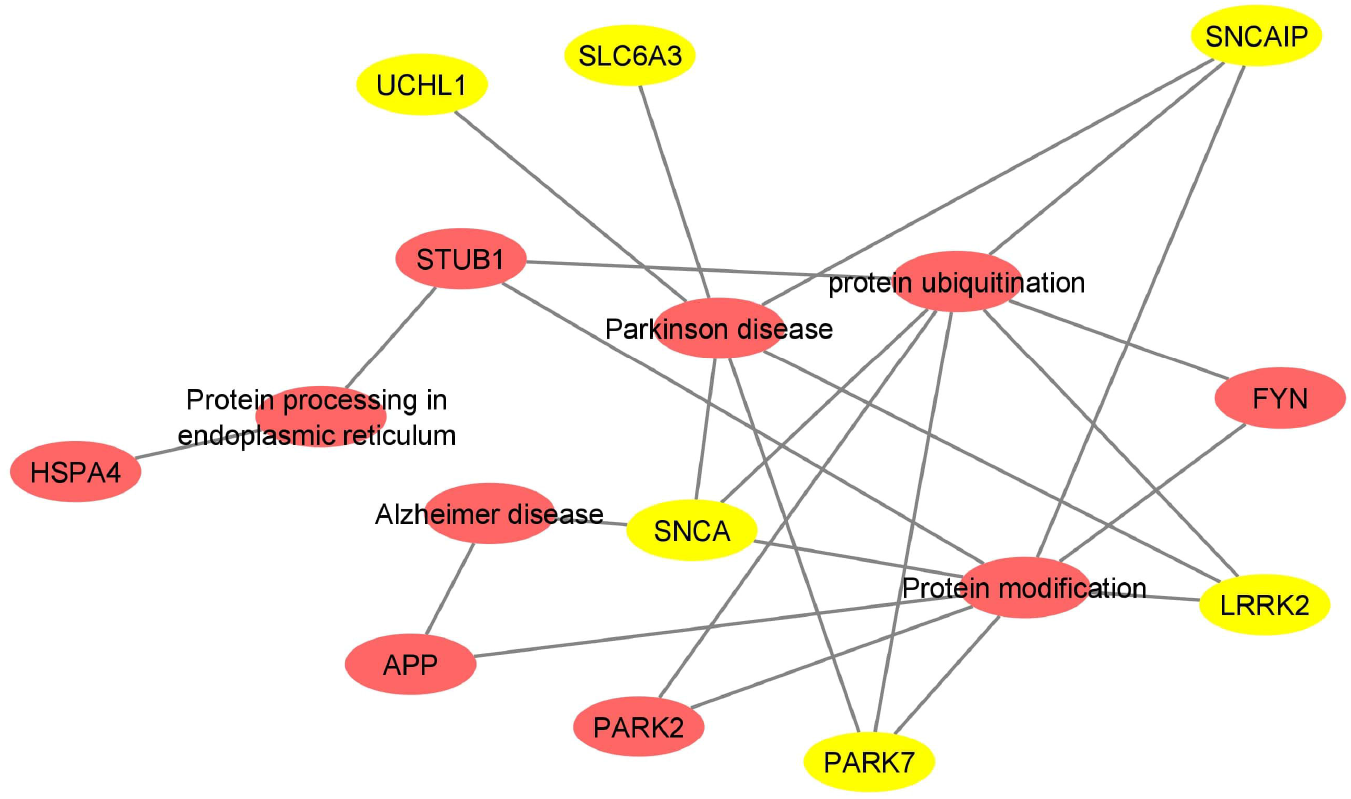
Network modelling of selected proteins on the basis of shared common biological pathway

### Retrieval of the herbal compound used in PD as a ligand and their drug likeness property

Retrieval of the twelve selected herbal compounds (Methylxanthine PubChem CID: 2519, Acanthopanax PubChem CID: 71307447, Alpinia epoxide PubChem CID: 100927652, Bacopaside1 PubChem CID: 10629555, Baicalein PubChem CID: 5281605, Chrysanthemum PubChem CID: 132503335, Clausena lansium PubChem CID: 11804919, Withania somnifera PubChem CID: 16760705, Tripterygium PubChem CID: 118701083, Resveratrol PubChem CID: 445154, Plumbagin PubChem CID: 10205, Pueraria glycoside PubChem CID: 44257229) was done using Pubchem compound database. The drug likeness properties of selected compound were calculated using Lipinski filter [23]. The Lipinski Rule of five is a thumb rule to evaluate drug likeness or determine if a chemical compound with a pharmacological or biological activity has properties that would make it an orally active drug in humans. Lipinski rule of 5 helps in distinguishing between drug like and non-drug like molecules. It predicts high probability of success or failure due to drug likeness for molecules complying with 2 or more of the following rules. Results of Lipinski filter for selected drug molecules following drug likeness properties are presented in (Table 2).

**Table 2.**
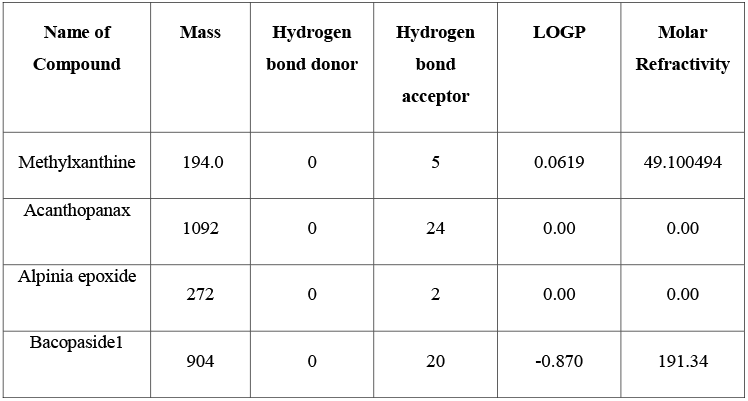
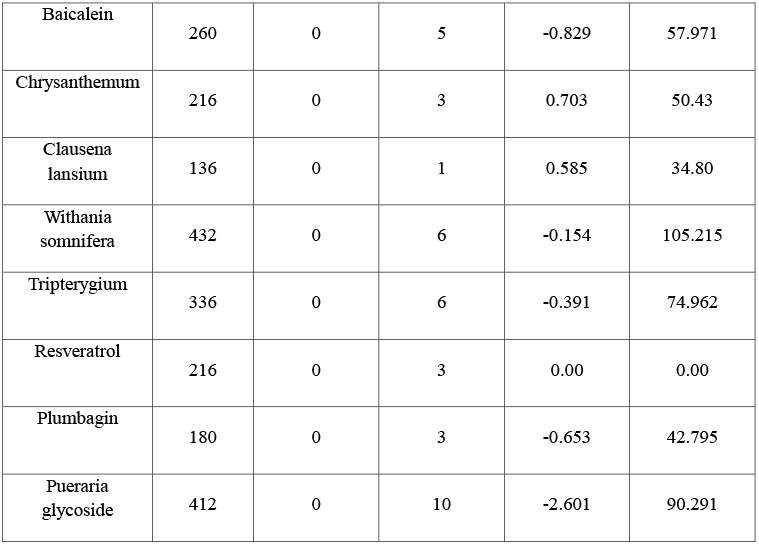
List of selected herbal compound which has been used as drug molecule for Parkinson disease and their drug likeness properties

### Molecular docking of twelve selected herbal compound with SNCA protein

Patchdock server was used for docking calculation. Before performing docking analysis active site prediction was done using Discovery studio 3.5. In PatchDock docking calculation, ligand molecules (Methylxanthine, Acanthopanax, Alpinia epoxide, Bacopaside1, Baicalein, Chrysanthemum, Clausena lansium, Withania somnifera, Tripterygium, Resveratrol, Plumbagin, and Pueraria glycoside molecules were used as drugs) interacted with prominent site of protein receptor identified using discovery studio3.5 with docking scores such as: 15942, 6390, 4254, 7212, 3786, 3608, 4820, 4140, 3692 and 4954 respectively (Table 3)(Fig 3). The receptor interacted residues were involved in interaction with selected herbal plants, which belongs to major active site of the SNCA protein, However, Methylxanthine compound was interacted with active site of SNCA with highly binding score of 15942, while the other herbal compounds that played their role as neuroprotective were interacted on the same position with low binding score in comparison to methylxanthine. As the methylxanthine compound was interacted with SNCA protein with more binding score as comparison to others, so they might be the prominent drug for PD (Table 3).

**Table 3.**
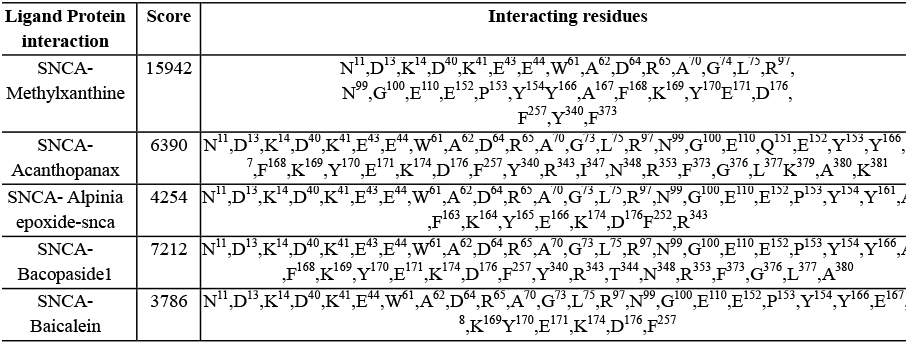
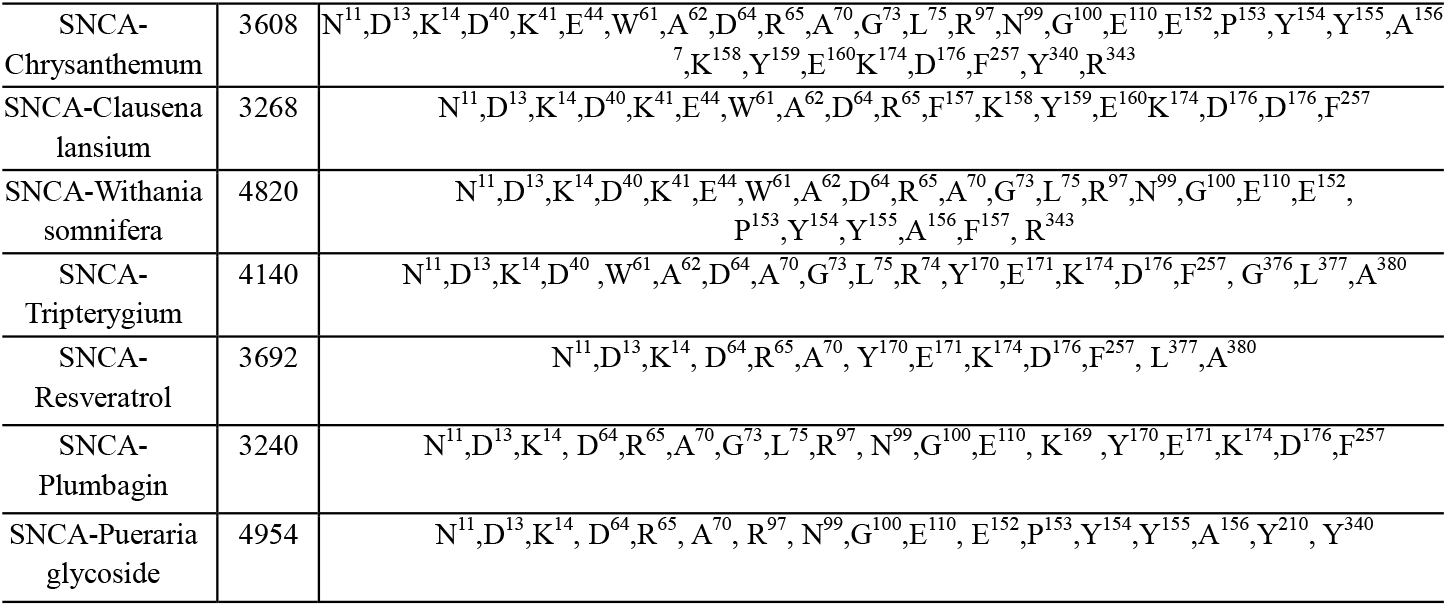
Molecular docking analysis of herbal compound which has been used as drug molecule for Parkinson disease and residues involved in interaction of SNCA protein with selected drug molecules

**Fig. 3.**
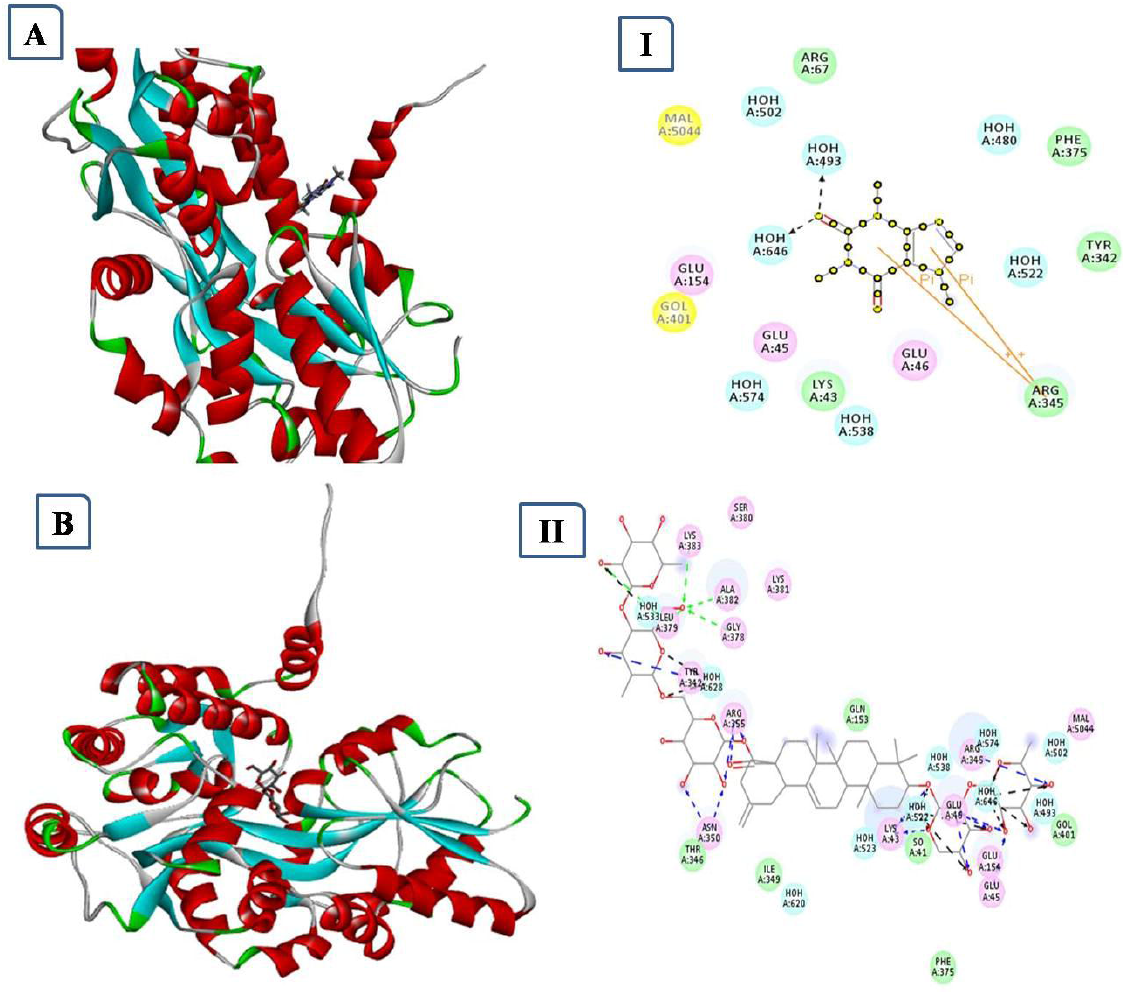
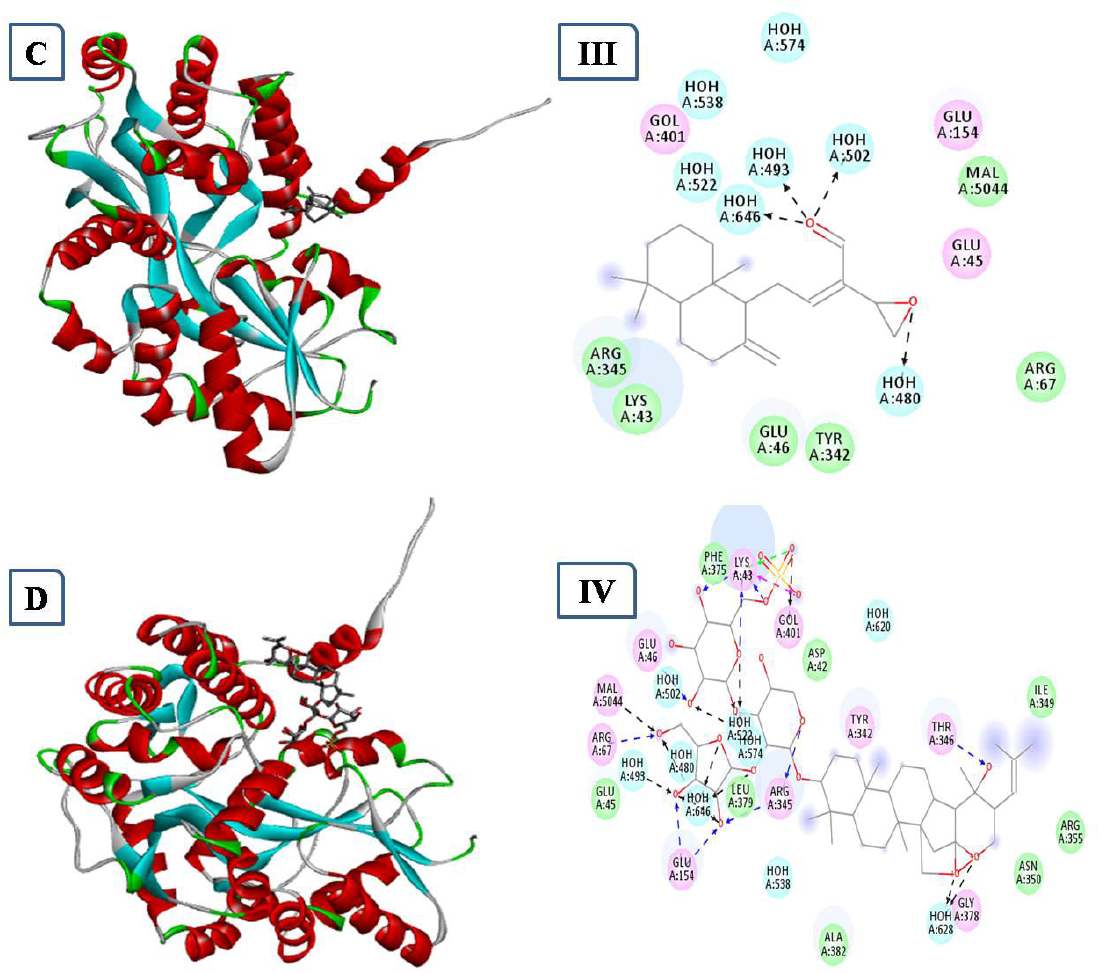
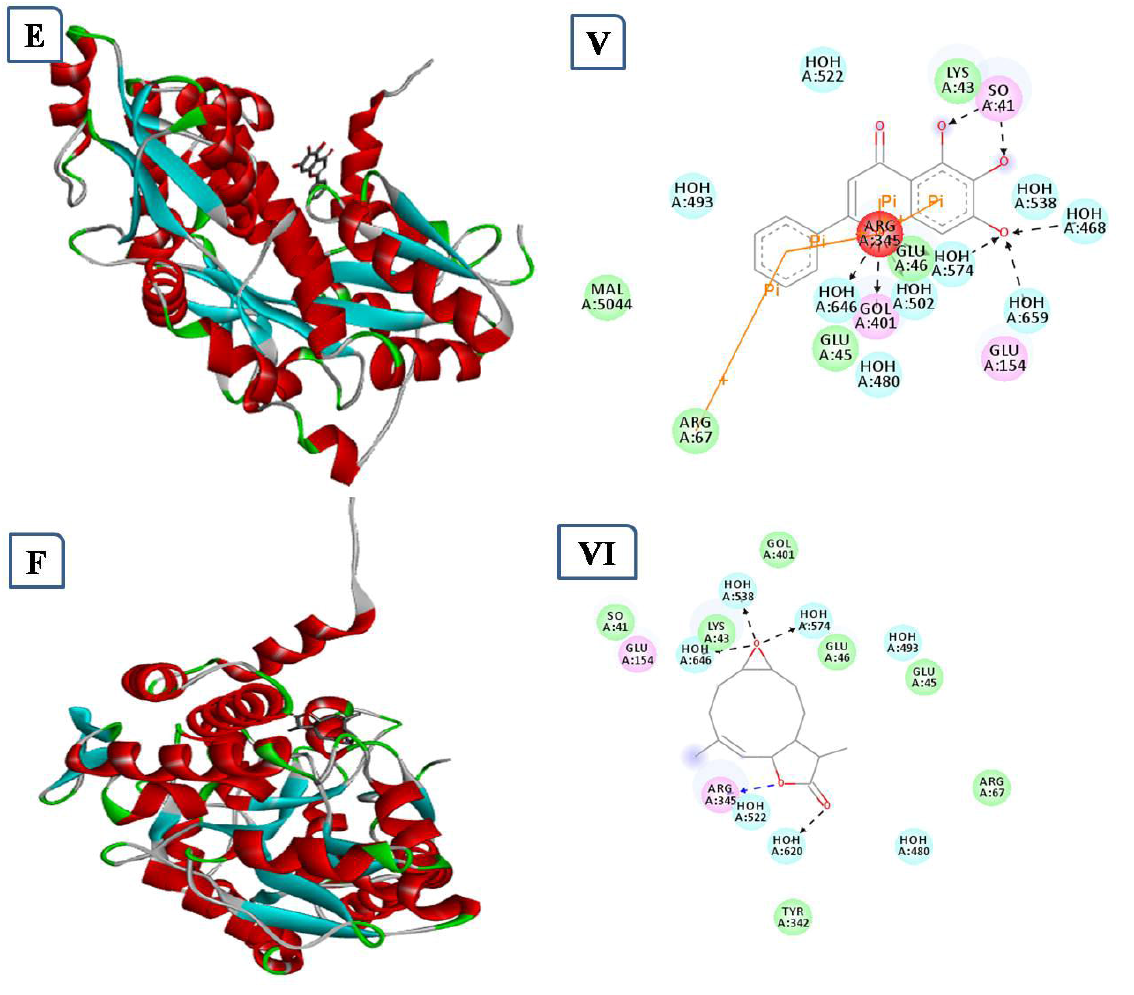
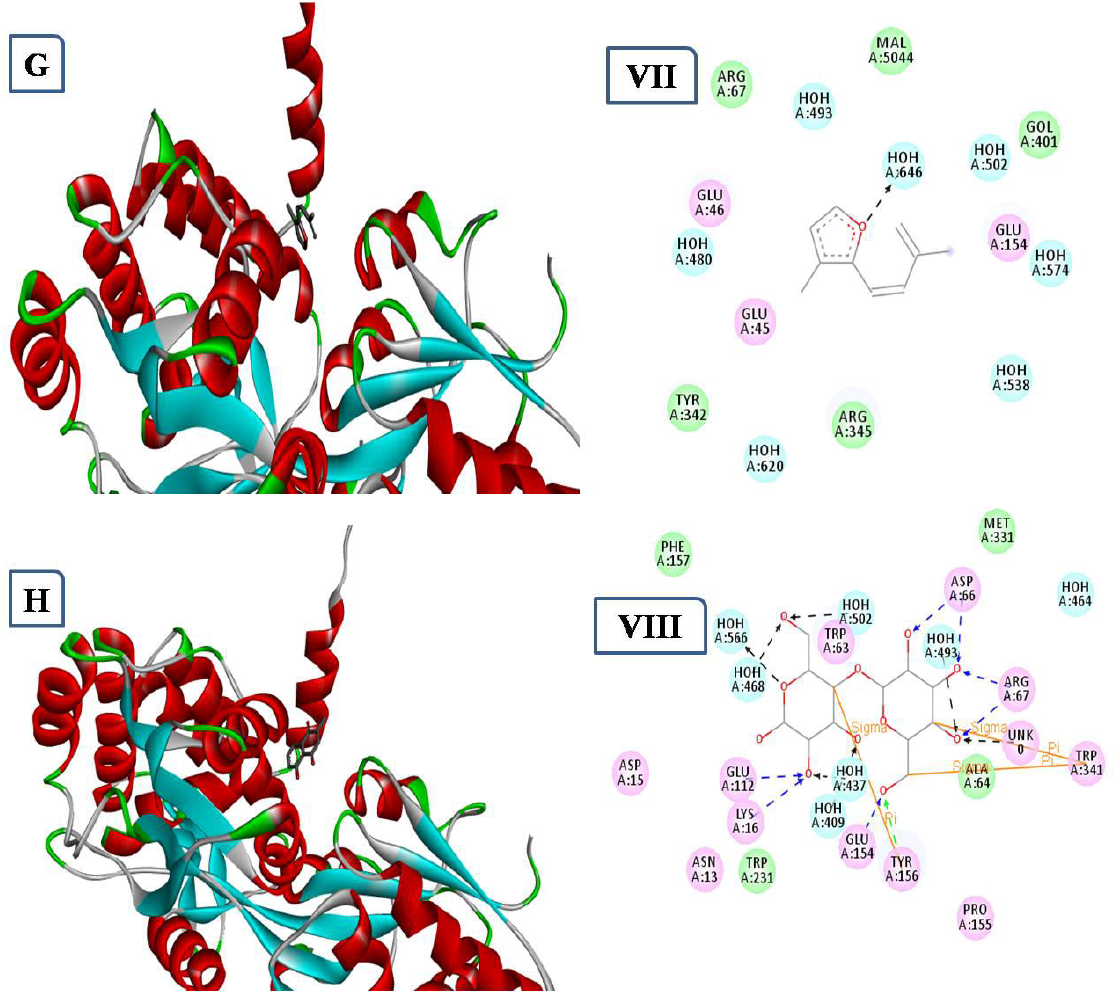
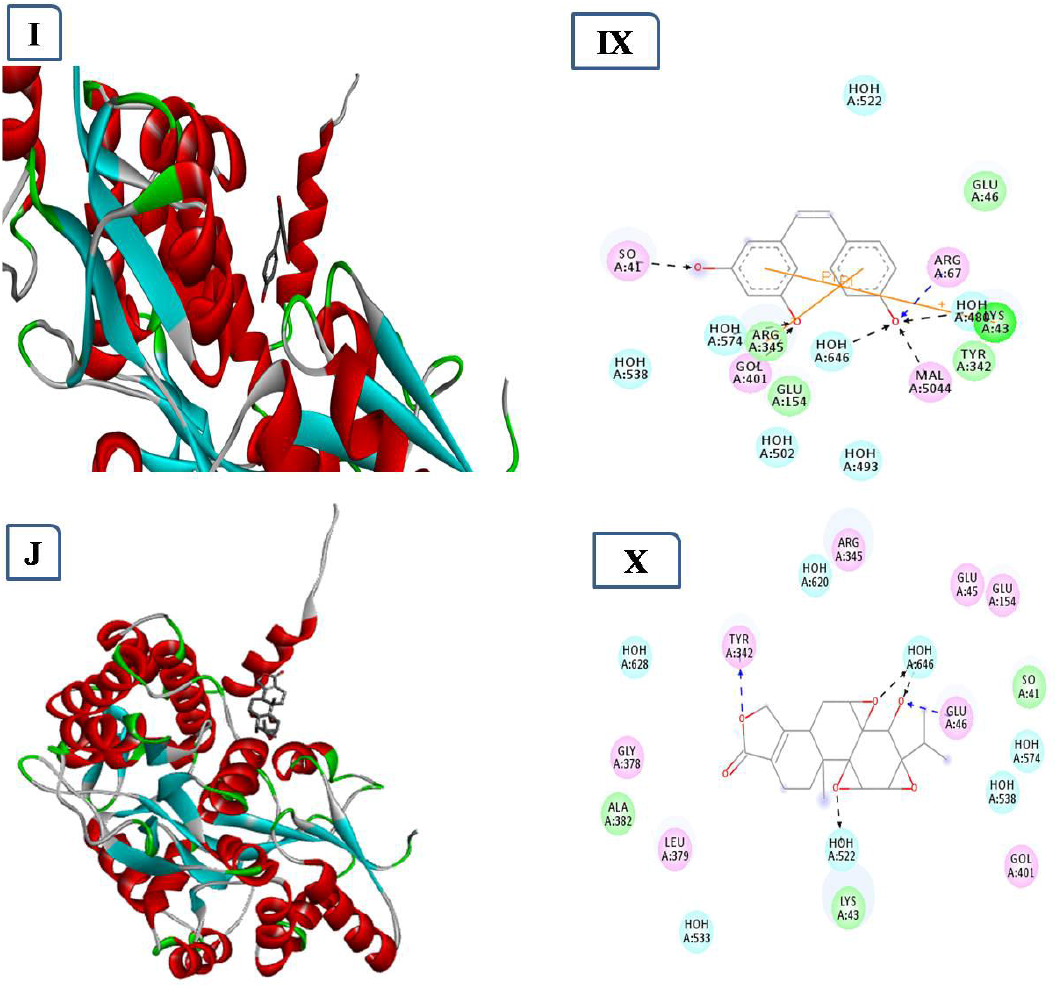
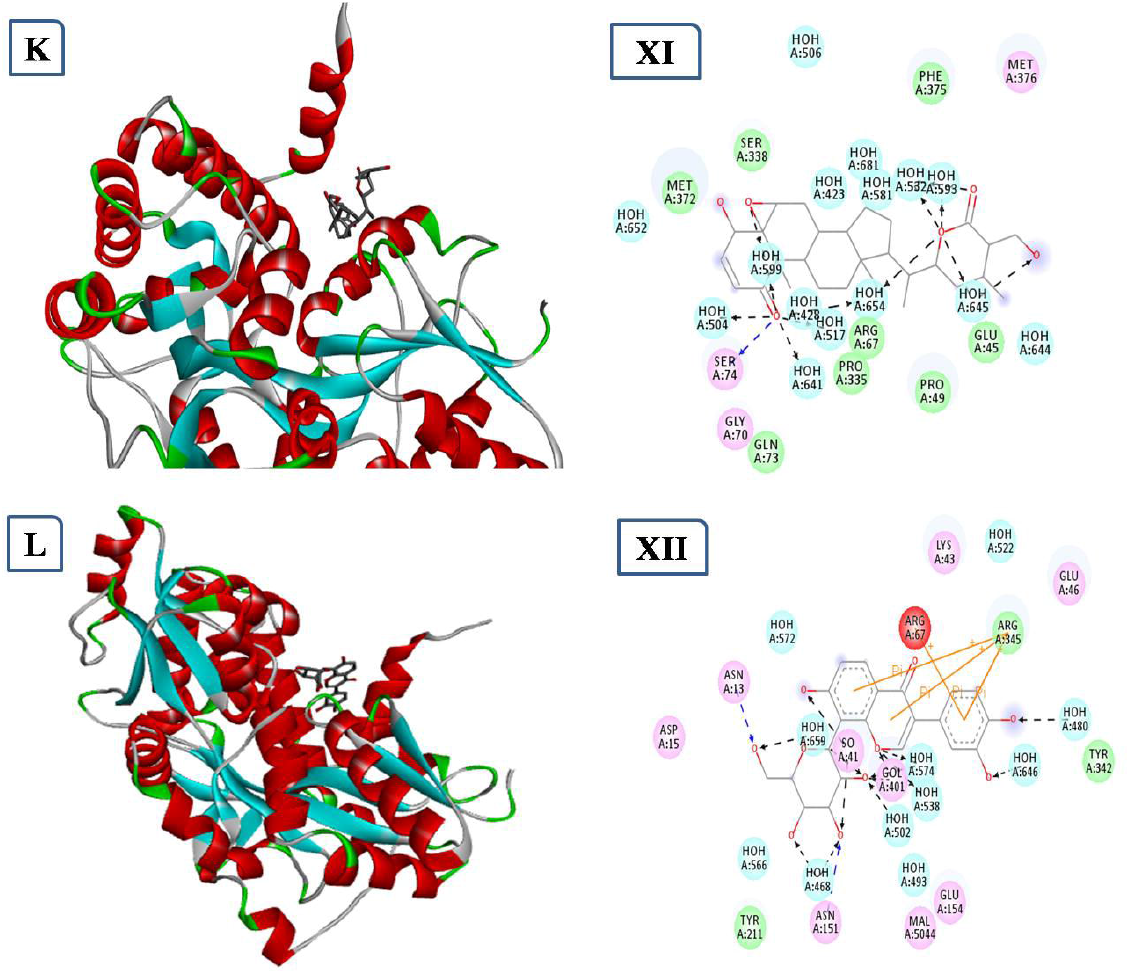
Interaction of selected herbal compounds with alpha synuclein protein and their interacting residues in 2D diagram, where interaction of selected herbal compounds Methylxanthine, Acanthopanax, Alpinia epoxide, Bacopaside1, Baicalein, Chrysanthemum, Clausena lansium, Withania somnifera, Tripterygium, Resveratrol, Plumbagin, Pueraria glycoside with SNCA protein were shown in images A, B, C, D, E, F, G, H, I, J, K, and L respectively. Whereas 2D interaction diagram of these compounds with SNCA were shown I, II, III, IV, V, VI, VII, VIII, IX, X, XI, and XII respectively.

## Discussion

Using the STRING database and cytoscape software, we studied the Interaction among proteins involved in neurodegeneration disease, primarily the Parkinson’s disease. In this networking all proteins were connected with each others. Further networking of selected proteins such as: APP, SNCA, STUB1, LRRK2, PARK1, SNCAIP, FYN, HSPA4, UCHL1, and SLC6A3 were performed on the basis of their shared common biological pathways. From this network, it has been found that the SNCA protein was more prominent interactor and involved in neurodegeneration to cause PD. Designing of drugs that can target the protein involved in neurodegeneration may be an effective approach for the treatment of PD [19]. Through computational molecular virtual screening of small molecules from natural compound MTX have been confirmed to directly inhibit these important proteins to prevent the PD, and were comparable with other natural compounds, such as: Acanthopanax, Alpinia epoxide, Bacopaside1, Baicalein, Chrysanthemum, Clausena lansium, Withania somnifera, Tripterygium, Resveratrol, Plumbagin, and Pueraria glycoside molecule which have been used as drugs in PD [26]. From this docking analysis we have found Methylxanthine as prominent drug molecule in comparison to other drugs to cure PD due to their higher binding efficiency (docking score: 15942).

## Conclusion

Parkinson’s disease (PD) is the second most common neurodegenerative disorder. Therefore the present study was aimed to identify the candidate protein based on its inhibitory effect on other proteins associated with neurodegeneration disease by using bioinformatics tools and algorithms. Although, there is no reliable treatment or drug to prevent Parkinson’s disease. However, our study has shown methylxanthine to be the potential target drugs that may plays important roles in the prevention of PD.

## List of abbreviations

GO: Gene Ontology
KEGG: Kyoto Encyclopedia of Genes and Genomes
PD: Parkinson’s disease
MTX: Methylxanthine

## Funding

This study was financially supported by IOE Scheme under Dev. Scheme No. 6031/2021).

## Authors’ contributions

Present study was designed and revised by NKR, NS, BK, AP and VNM. NKR and NS combinedly performed the bioinformatics analysis while the docking analysis was performed by NS. The manuscript was written and drafted by NKR and VNM. All authors have read and approved the manuscript.

## Acknowledgements

The authors kindly acknowledge Interdisciplinary School of Life Sciences (ISLS), and Information Centre for Bioinformatics, School of Biotechnology, Institute of Science, Banaras Hindu University for providing software and the computational facilities for bioinformatics analysis. We are highly grateful to the Department of Neurology (IOE), Institute of Science, Banaras Hindu University, Varanasi for extending laboratory facilities

## Authors’ information

Nishant Kumar Rana, Vijay Nath Mishra & Abhishek Pathak, Department of Neurology, Institute of Medical Sciences, Banaras Hindu University Varanasi-221005, U. P., India

Neha Srivastava, Department of Bioinformatics, Mahila Maha Vidyalaya, Banaras Hindu University, Varanasi-221005, U. P., India

Bhupendra Kumar, Department of Zoology, Institute of Science, Banaras Hindu University, Varanasi-221005, U. P., India

## Declaration of competing interests

The authors declare that they have no competing interests.

## Ethics approval and consent to participate

No ethics approval was required for the study.

## Consent for publication

All the authors have agreed for the publication without any conflict of interest.

## Notes

### Competing Interest Statement

The authors have declared no competing interest.

